# Extracellular Matrix Hydrogels Promote Expression of Muscle-Tendon Junction Proteins

**DOI:** 10.1101/2021.03.24.436805

**Authors:** Lewis S. Gaffney, Zachary G. Davis, Camilo Mora-Navarro, Matthew B. Fisher, Donald O. Freytes

## Abstract

Muscle and tendon injuries are prevalent and range from minor sprains and strains to traumatic, debilitating injuries. However, the interactions between these tissues during injury and recovery remain unclear. Three-dimensional tissue models that incorporate both tissues and a physiologically relevant junction between muscle and tendon may help understand how the two tissues interact. Here, we use tissue specific extracellular matrix (ECM) derived from muscle and tendon to determine how cells of each tissue interact with the microenvironment of the opposite tissue resulting in junction specific features. ECM materials were derived from the Achilles tendon and gastrocnemius muscle, decellularized, and processed to form tissue specific pre-hydrogel digests. ECM materials were unique in respect to protein composition and included many types of ECM proteins, not just collagens. After digestion and gelation, ECM hydrogels had similar complex viscosities which were less than type I collagen hydrogels at the same concentration. C2C12 myoblasts and tendon fibroblasts were cultured in tissuespecific ECM conditioned media or encapsulated in tissue-specific ECM hydrogels to determine cell-matrix interactions and the effects on a muscle-tendon junction marker, paxillin. ECM conditioned media had only a minor effect on upregulation of paxillin in cells cultured in monolayer. However, cells cultured within ECM hydrogels had 50-70% higher paxillin expression than cells cultured in type I collagen hydrogels. Contraction of the ECM hydrogels varied by the type of ECM used. Subsequent experiments with varying density of type I collagen (and thus contraction) showed no correlation between paxillin expression and the amount of gel contraction, suggesting that a constituent of the ECM was the driver of paxillin expression in the ECM hydrogels. In addition, the extracellular matrix protein type XXII collagen had similar expression patterns as paxillin, with smaller effect sizes. Using tissue specific ECM allowed for the de-construction of the cell-matrix interactions similar to muscletendon junctions to study the expression of MTJ specific proteins.

**Impact Statement:** The muscle-tendon junction is an important feature of muscle-tendon units; however, despite crosstalk between the two tissue types, it is overlooked in current research. Deconstructing the cell-matrix interactions will provide the opportunity to study significant junction specific features and markers that should be included in tissue models of the muscletendon unit, while gaining a deeper understanding of the natural junction. This research aims to inform future methods to engineer a more relevant multi-tissue platform to study the muscletendon unit.

## 1. Introduction

Musculoskeletal injuries involving muscle and tendon are prevalent and range from minor sprains and strains to debilitating injuries from volumetric muscle loss or tendon rupture.^1^ In addition to acute injuries, muscle and tendon disorders can be caused by aging, dystrophies and inflammation.^2^ Muscle and tendon tissue are inherently capable of healing after minor injuries, but in the case of massive defects or disease, loss of function can occur. Regenerative medicine is a promising new area to combat loss of function, and greatly improve the quality of life for those affected. Currently researchers in regenerative medicine have made great improvements in single tissue engineering constructs, but there are far fewer engineered tissue systems that incorporate interfaces, including the muscle-tendon junction.

A crucial part of the muscle-tendon unit is the myotendinous junction, which transmits the contractile force of the muscle through the tendon to the skeletal system (Figure 1). At the myotendinous junction, muscle cells anchor to tendon matrix through the integrin-mediated complex.^3^ The integrin-mediated complex is made up of several intracellular proteins: paxillin, vinculin, tensin and talin, that bind actin within the muscle cells to the β1 subunit of the myotendinous junction specific α7β1 integrin.^3^ In tendon fibroblasts, paxillin is an important component for cell adhesion, motility and is linked to focal adhesion turnover, all of which are important factors in regeneration of tendon.^4^ As tendon fibroblasts mature, paxillin localization in the cell membrane decreases along with other focal adhesion proteins leading to decreased cell motility which has implications for healing in mature tendon.^5^ Paxillin expression in muscle cells occurs at the junction, where the sarcolemma binds to tendon ECM forming numerous focal adhesion complexes that bind to α7β1 integrins in response to mechanical stimulation.^3,6^ The specialized attachment (not present throughout the rest of the muscle tissue) is an example of specific structural component for muscle cells present at the myotendinous junction that could play a vital role in the function of the muscle-tendon unit. Figure 1C shows where muscle fibers meet tendon ECM indicated by the white dashed line and arrows, and figure 1D shows increased paxillin localization, throughout this area. In addition to the integrin mediated complex, type XXII collagen is a junction specific extracellular matrix protein produced by muscle and tendon cells.^7,8^ Type XXII collagen is a basement membrane protein present at tissue junctions, especially the myotendinous junction contributing to mechanical stability of the MTJ.^8,9^ In studies of proteomic analysis of muscle-tendon units, type XXII collagen was isolated in the myotendinous region and is thought to interact with α7β1 integrins.^10,11^ While there is some understanding of the cellular response to various disorders in muscle and tendon tissue individually, there is less understanding of how disorders in a single tissue may further inhibit reparative processes and decrease function of the muscle-tendon unit.^2^ Currently, there are a limited number of techniques that aim to re-capitulate the multi-phasic structure of the MTJ, including the native interface between the tissues. The first multi-phasic tissues to model the muscle-tendon unit were engineered by seeding muscle cells onto a fibroblast monolayer, creating an inner muscle tissue and outer tendon-like tissue which expressed MTJ specific proteins.^12^ These constructs expressed paxillin, although the protein expression was not localized to the interface of the cells. More recently 3D printing with bio-inks and polycaprolactone has been used to create an interface between fibroblasts and muscle cells.^13,14^ 3D printing was successful in spatially organizing the two cell types and allowing for differentiation of the cells to form multi-phasic muscle-tendon constructs. However, these systems relied on cell-cell interactions to produce MTJ expression and did not incorporate the extracellular matrix (ECM) microenvironment that exists within the muscle-tendon unit, which prevented the ability to analyze how specific cells at the junction interact with the environment of the opposite tissue, resulting in junction specific phenotypes and proteins.

**Figure 1:**
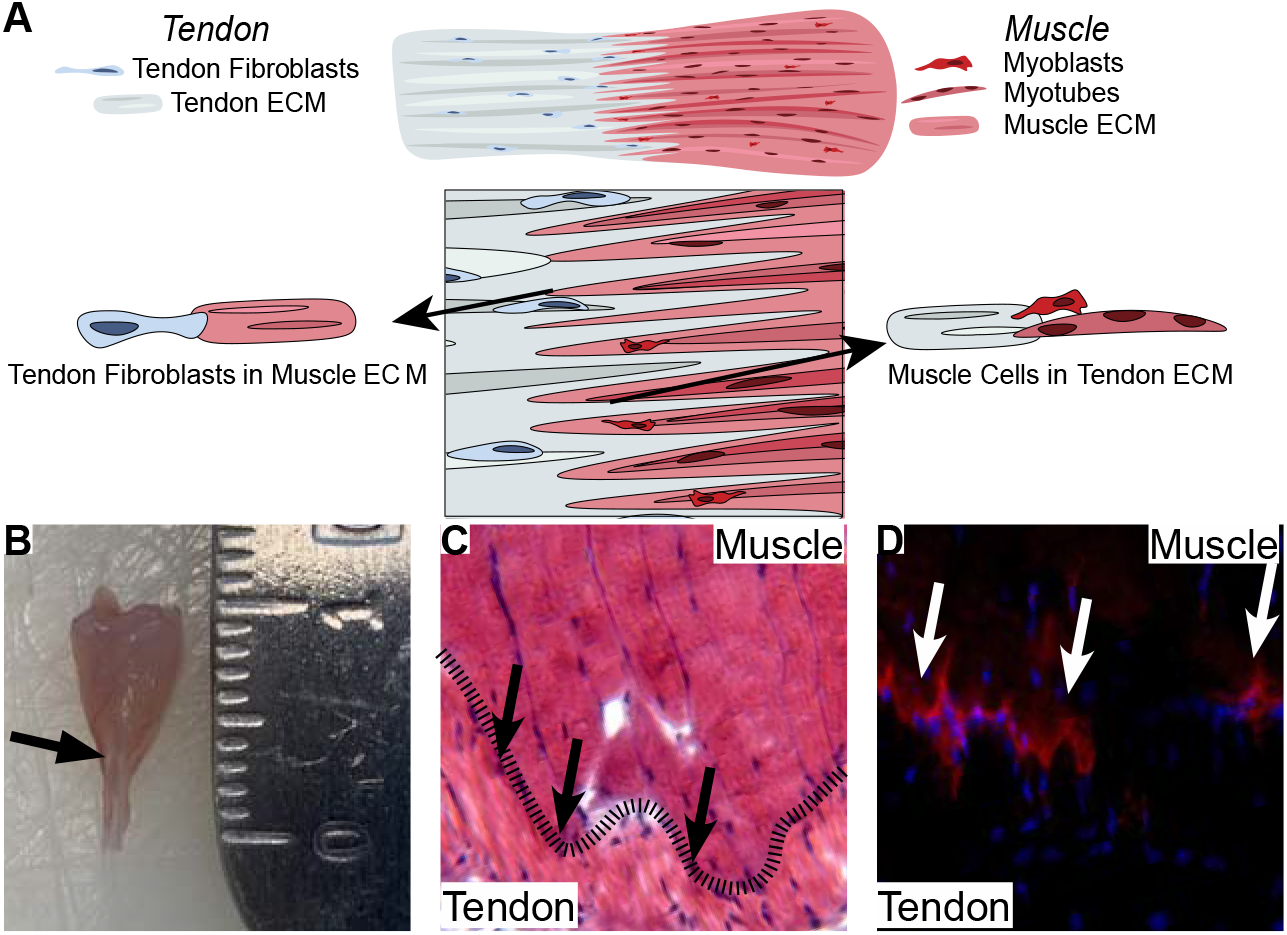
ECM interactions and paxillin expression at the myotendinous junction. A) Graphical description of ECM-cell interactions at the myotendinous junction. Where muscle and tendon meet, cells within one tissue interact with ECM of the other tissue. This interaction could be responsible for production of MTJ specific proteins such as paxillin, which is part of the integrin mediated complex attaching muscle cells to tendon ECM. B) A mouse muscle tendon unit (Achilles tendon and gastrocnemius muscle) dissected from mouse hindlimb was sectioned for C) hematoxylin and eosin staining, the dashed line indicating the junction of muscle and tendon, D) anti-paxillin immunohistochemical staining. Arrows in C and in D indicate the junction of tendon (bottom) and muscle (top). In D, at the junction there is localization of paxillin expression in the ends of large muscle fibers, where the sarcolemma interacts and binds to tendon ECM through the accumulation of focal adhesion complexes containing paxillin.

Controlling cell interactions with the ECM microenvironment of the MTJ may allow localization of protein expression and formation of native-like focal adhesion in cell construct models. Beyond mechanical cues, ECM materials derived from tissues contain tissue-specific factors after decellularization and processing, which can influence cell behaviors.^15,16^ ECM materials contain tissue specific composition of ECM proteins and growth factors contribute to inherent bioactivity.^17^ This tissue specific composition has been known to influence cell phenotypes such that stem cells or progenitor cells differentiate along similar pathways to the source tissues.^17^ Examples include decellularized bone^18^, muscle^19–22^ and tendon^23–25^, where in each case, ECM increased expression of tissue specific cell phenotypes *in vitro* or functionality of the tissues after healing *in vivo*. More recently, decellularized muscle-tendon units were used *in vivo* and *in vitro* to increase myogenic expression in satellite cells and improve function in a muscle defect model compared to a collagen implant alone.^26^ However, such models do not allow us to determine how cells interact with neighboring tissue ECM.

The objective of this research is to determine if tissue specific ECM can promote expression of MTJ integrin associated proteins as well as ECM proteins specific to the junction, in myoblast and fibroblast cells. Using ECM materials derived from muscle and tendon allows for the culture of muscle and tendon cells in a three-dimensional microenvironment similar to a muscle-tendon unit. ECM materials can provide an extra component of signaling to cells cultured within the tissues that goes beyond cell-cell interactions. Incorporating a microenvironment similar to the native tissue, where muscle cells are in contact with tendon ECM and tendon cells in contact with muscle ECM could add function to *in vitro* models that have muscle and tendon components. Here, we present methods to de-construct the myotendinous junction by using tissue specific ECM with different cells to study how cells interact with the opposite tissue’s microenvironment.

## 2. Methods

### 2.1 Muscle and Tendon derived ECM Hydrogels

Extracellular matrix materials were derived from porcine tissue harvested from 3-18 month old pigs (NCSU Swine Education Unit, Raleigh, NC) after sacrifice for other IACUC approved experiments ongoing in our lab. To obtain specific tissues, gastrocnemius muscle and Achilles tendon were excised as a muscle-tendon unit from the lower shank of the pigs. Achilles tendon was dissected distal to the myotendinous junction and cleaned of connective tissue. Gastrocnemius was separated from the tendon proximal to the myotendinous junction and cleaned of connective tissue. (Figure 2A, upper left) Both muscle and tendon tissues were then washed and sliced into patches less than 1mm thick and frozen at −80°C.

**Figure 2:**
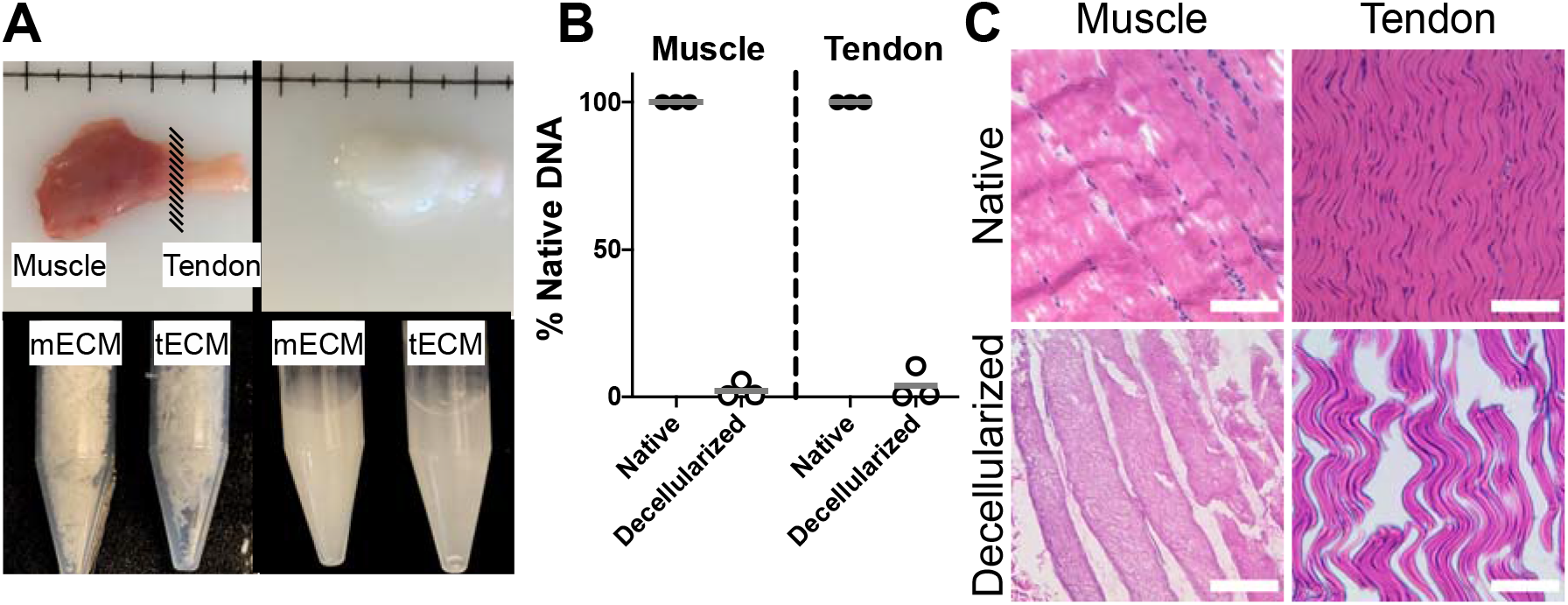
Decellularization methods of separate muscle and tendon. A) Porcine muscle tendon unit was excised from hindlimb, upper left, (shown still connected, for separated materials, muscle and tendon were separated) and decellularized, upper right. Tissues were lyophilized and ground into powder, lower left, and digested with pepsin, lower right. B) Presence of DNA was measured with PicoGreen assays. DNA content in native tissue was used to normalize %DNA removal. Muscle and tendon had a mean of 98% and 96% less DNA than native tissue after decellularization, respectively (n=3, mean represented by bar). C) Hematoxylin and eosin stains before and after decellularization also confirms removal of nuclear material from muscle and tendon. Scale bars are 200 μm.

Tissues were decellularized with adaptations from a previously described method^22^. Muscle and tendon tissue were washed separately in 50ml conical tubes with the protocol detailed in Table 1. Decellularized tissues were then lyophilized then ground in a mill to yield a powder (Figure 2A, lower left). Tissue specific hydrogels for muscle and tendon (mECM and t-ECM) were produced from the powder by digesting with 20:1 weight ratio of pepsin from porcine submucosa (Sigma-Aldrich) in .01 M HCl at a concentration of 10mg/ml. ECM digests (Figure 2A, lower right) were neutralized and buffered, m-ECM and t-ECM self-assembled to form stable hydrogels after 45 minutes at 37°C. ECM hydrogels were formed, cryosectioned and stained with DAPI, anti-Paxillin and anti-Col22a1 to verify removal of nuclear content and absence of proteins of interest (Supplemental Figure 1).

**Table 1:** Muscle and Tendon Decellularization Protocol. Decellularization protocol details for muscle and tendon tissue based on protocol from Wolf et. al.^22^ Tendon and muscle tissues were washed in separate 50 ml conical tubes in each of the solutions for the detailed amount of time.

| Solution | Time |
| --- | --- |
| 0.2%Trypsin/EDTA | 2 hrs. |
| DI Water | 30 min. |
| 2X PBS | 30 min. |
| 0.5% Sodium Deoxycholate | 5 hrs. |
| DI Water | 30 min. |
| 2X PBS | 30 min. |
| 0.5% Sodium Deoxycholate | 14-16 hrs. |
| 1.0%Triton X-100 | 1 hr. |
| DI Water | 30 min. |
| 0.1% Peracetic Acid/4.0% Ethanol | 2 hrs. |
| 1X PBS | 30 min. (2X) |
| Di Water | 30 min. (2X) |

### 2.2 Proteomics of ECM Derived Materials

#### Sample Preparation

Decellularized mECM and tECM powder was pooled from multiple animals. Discovery proteomic workflow was used to characterize and compare the overall protein composition of decellularized biomaterials. To generate adequate data for analysis ~10 mg of mECM or tECM was used. Samples were suspended in 1 mL of 50 mM ammonium bicarbonate (pH = 8.0) with 5% w/v sodium deoxycholate (SigmaAldrich) for digestion and homogenization. Samples were homogenized using an OMNI tissue homogenizer and a Fisher Scientific Sonic Dismembrator Model 120. Samples were then centrifuged at 10,000 x g for 10 min, and a bicinchoninic acid assay (BCA assay) was performed on the supernatant. Based on this BCA measurement, samples were normalized to 50 ug of protein and a filter-aided sample preparation (FASP) protocol was employed to digest and clean the samples prior to mass spectrometry analysis.^27^

#### Liquid Chromatography-Mass Spectrometry (LC-MS)

All samples were run using a Thermo Scientific Easy Nano-LC 1200 with an EASY-Spray source complexed to a ThermoFisher Scientific Orbitrap Exploris 480. A ThermoFisher Scientific Acclaim PepMap 100 trap column (C18 LC Columns, 3 μm particle size, 75 μm ID, 20 mm length) was utilized in line with an EASY-Spray analytical column (2 μm particle size, 75 μm ID, 250 mm length) at 35 °C. A 140 minute gradient using solvents A: 2% acetonitrile (ACN), 0.1% formic acid (FA) (aq) and B: 20% H2O, 0.1% FA, 79.9% ACN was run using the following steps at 300 nL/minute: 0 min - 5% B, 2 min – 5 % B, 107 min – 25% B, 122 min – 40% B, 123 min – 95% B, 140 min – 95% B. The mass spectrometer was run in data dependent acquisition (DDA) mode with a cycle time of 2 seconds. 2 μL of volume was injected for each sample, containing approximately 1 μg of protein material total.

#### Data Analysis

Raw data were loaded into Proteome Discoverer (PD) 2.4.0.305 (ThermoFisher Scientific) for analysis. A label free quantitation (LFQ) workflow was used, and the data was normalized by total peptide amount using PD. For peptide searching, the protein FASTA database was downloaded via Proteome Discoverer from SwissProt (fully annotated) and TrEMBL (unreviewed proteins) databases for Sus Scrofa (taxonomy ID = 9823) (Version 2017_09, Released October 25, 2017). A maximum of 2 equal mods were used per peptide. The following post translational modifications (PTMs) were accommodated in the search algorithm (modified amino acids in parentheses): oxidation (M), deamidation (N, Q). Gene Ontology (GO) functional classification analysis (protein class) was performed using the “Gene List Analysis” tool from www.pantherdb.org using Sus Scrofa as the organism and plots were constructed via PRISM or BioVenn (Figure 3).^28^ Associated ECM proteins and subunits were listed in according to Naba et. Al (Figure 3).^29^ Proteomics data has been uploaded to a publicly available repository, https://github.com/Translational-Orthopaedic-Research-Lab/Muscle_Tendon_ECM_Material_Proteomics.

**Figure 3:**
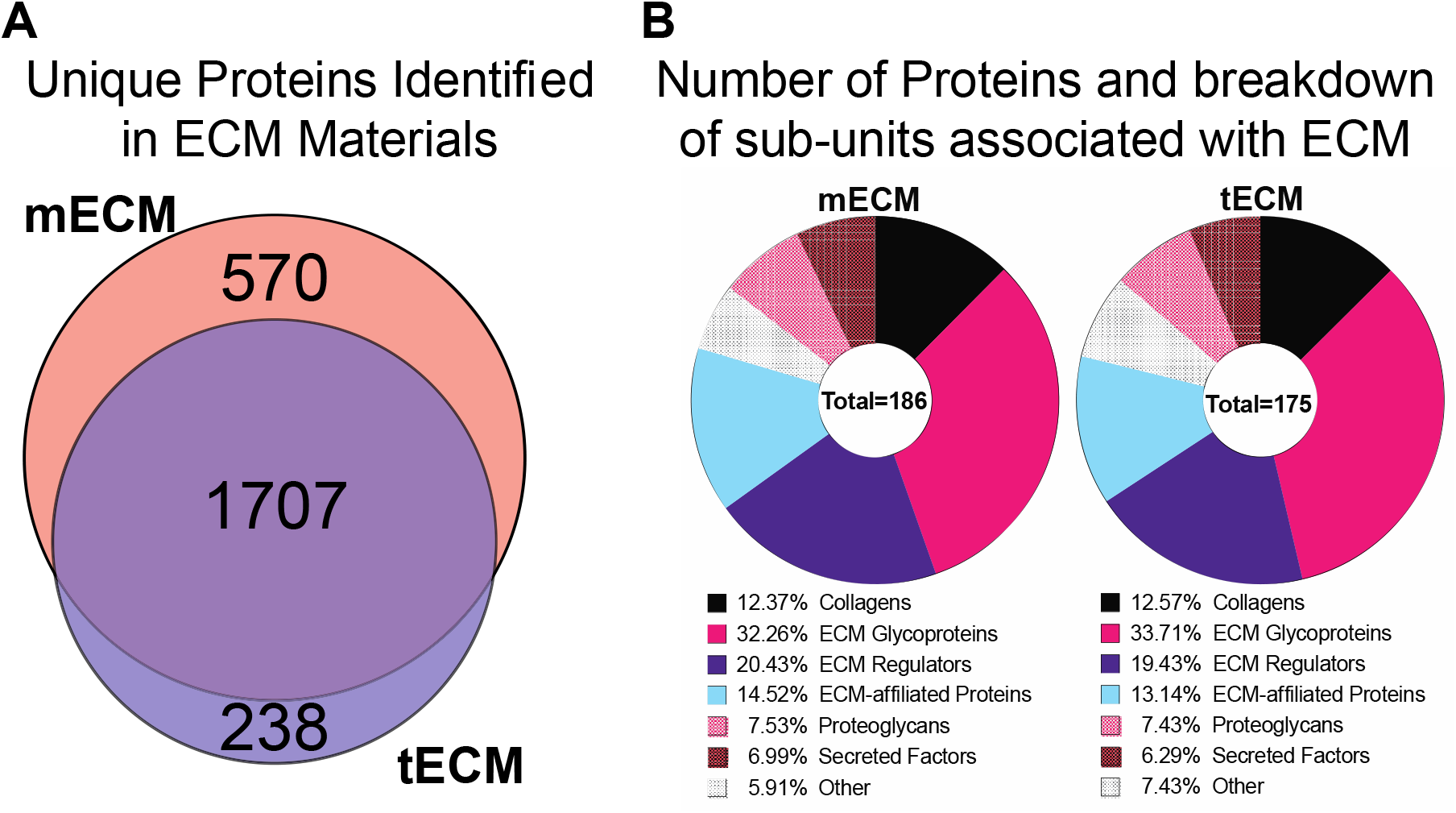
Discovery proteomics analysis of muscle and tendon derived ECM. A) Venn diagram showing the overall number of proteins identified per biomaterial and the common matches. mECM and tECM had 570 and 238 unique proteins identified in each of the materials. B) The number of proteins identified as ECM associated proteins was very similar, 186 protiens for mECM and 175 proteins for tECM. Pie charts show the percentage of total ECM associated proteins identified and classified for each biomaterial.

### 2.3 Rheological Properties of Extracellular Matrix Hydrogels

Rheological measurements were taken using a MCR 92 (AntonPaar, Graz, Austria) and following the parameters established by Freytes et al.^30^ Oscillatory shear strain was determined to be the best technique as it would prevent destruction of the self-assembly while allowing for the collection of complex viscosity. 5% strain was used based on prior work with similar ECM hydrogels.^30^ Complex viscosity was determined to be optimal as a higher complex viscosity would show a greater resistance to movement of the hydrogel, while a lower complex viscosity would show more flow was possible with applied forces-as would come from cells within the hydrogel system. Briefly, 5mg/mL hydrogels of type I collagen, tECM and mECM and 2mg/mL hydrogels of type I collagen were pipetted onto the bottom plate of the rheometer after neutralization and buffering. The 25mm parallel measuring plate (Anton Paar, Graz, Austria) was lowered onto the solution and excess material was trimmed away. The lower plate was heated to 37°C and left for 45 minutes to allow for full self-assembly of the hydrogel prior to initiating a frequency sweep from 100 rad/s to 25 rad/s at a constant 5% oscillating shear strain. n=3 samples from each material were tested. Data is presented as the mean, with the SD shaded (Figure 4).

**Figure 4:**
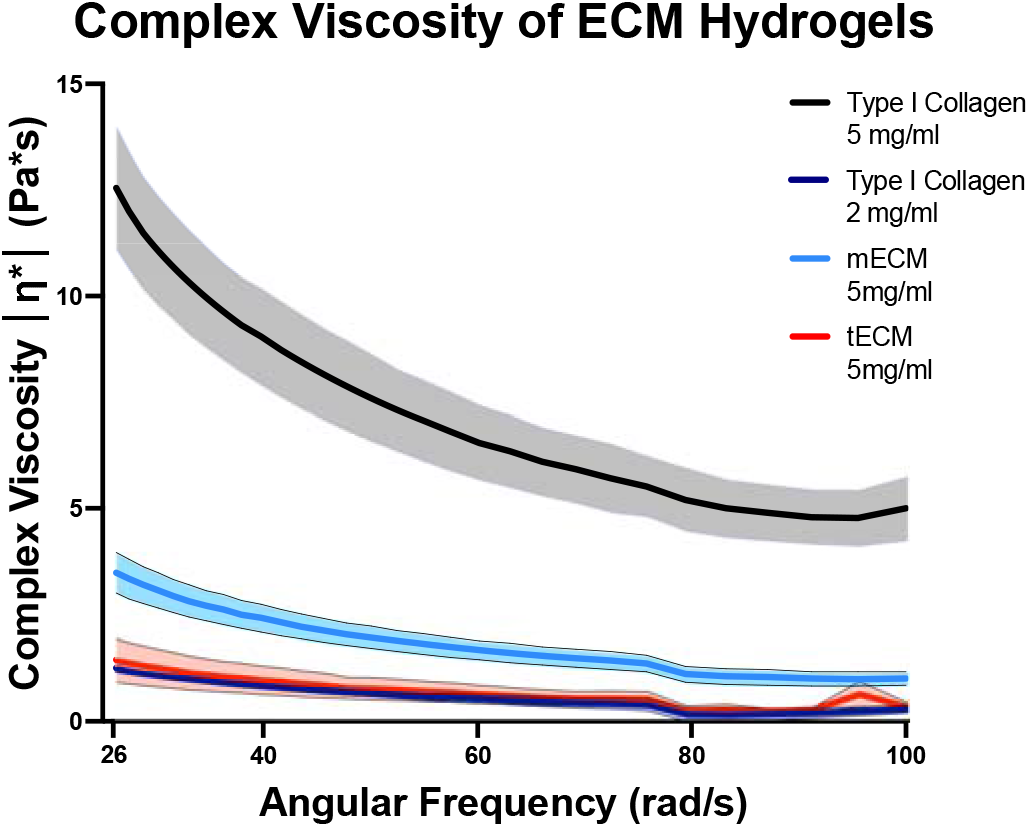
ECM hydrogels were measured with a rheometer to determine complex viscosity across a range of 26-100 rad/s oscillations to 5% shear strain. Type I collagen at 5 mg/ml was the most viscous material of the 4 measured across the oscillatory range, which could have an impact on cell’s ability to contract the tissue. ECM hydrogels and type I collagen at 2 mg/ml had similar rheological properties. Data is presented as mean, and standard deviation for n=3 samples per condition.

### 2.4 Cell Isolation and Preparation

C2C12 mouse myoblast cells were obtained from ATCC (C2C12s). Mouse tendon fibroblasts were isolated from mouse Achilles tendon (TFs). Briefly, hindlimbs were removed from C57BL/6 mice after sacrifice for another IACUC approved experiment. The Achilles tendon was excised from the limbs using sterile procedures in a biosafety cabinet. After excision, tendons were placed in a 6 well plate, minced with a scalpel, and left to soak in proliferation media, described below. After 24 hours, tendon tissues were removed to avoid contamination. Both cell-lines were cultured in proliferation media: Dulbecco’s Modified Eagle Media, DMEM (Thermofisher), 10% FBS (Genesee Scientific) and 1% Penicillin Streptomycin (Thermofisher). Cells were used for experiments between passage 3 and passage 10 (within previously used ranges for C2C12s and 3T3 fibroblast cells).^22^

### 2.5 Presence of MTJ Expression in Cells Cultured with ECM Conditioned Media

C2C12s were seeded on cell culture plates and allowed to attach overnight. 400ug/ml of either type I collagen, mECM or tECM was added to supplement media (Figure 5A). 400ug ECM/ml of media has been previously described to influence macrophage phenotypes^31^ and was the limit of ECM concentration before gelation of media started to occur. Cells were lysed on day 5 for gene expression analysis. 3 separate experimental replicates of 2 individual wells, combined after culture, were analyzed for gene expression analysis. RNA was isolated using EZNA RNA isolation kits (Omega Bio-Tek). PCR analysis was done using SYBR Green (ThermoFisher) and primers for GAPDH (forward: AGGTCGGTGTGAACGGATTTG, reverse: GGGGTCGTTGATGGCAACA) and Paxillin (forward: AACGGCCAGTGTTCTTGTCAG, reverse: CACCGCAATCTCCTGGTATGT). Paxillin gene expression was determined with q-PCR, and ΔCT was normalized to housekeeping gene, GAPDH.

**Figure 5:**
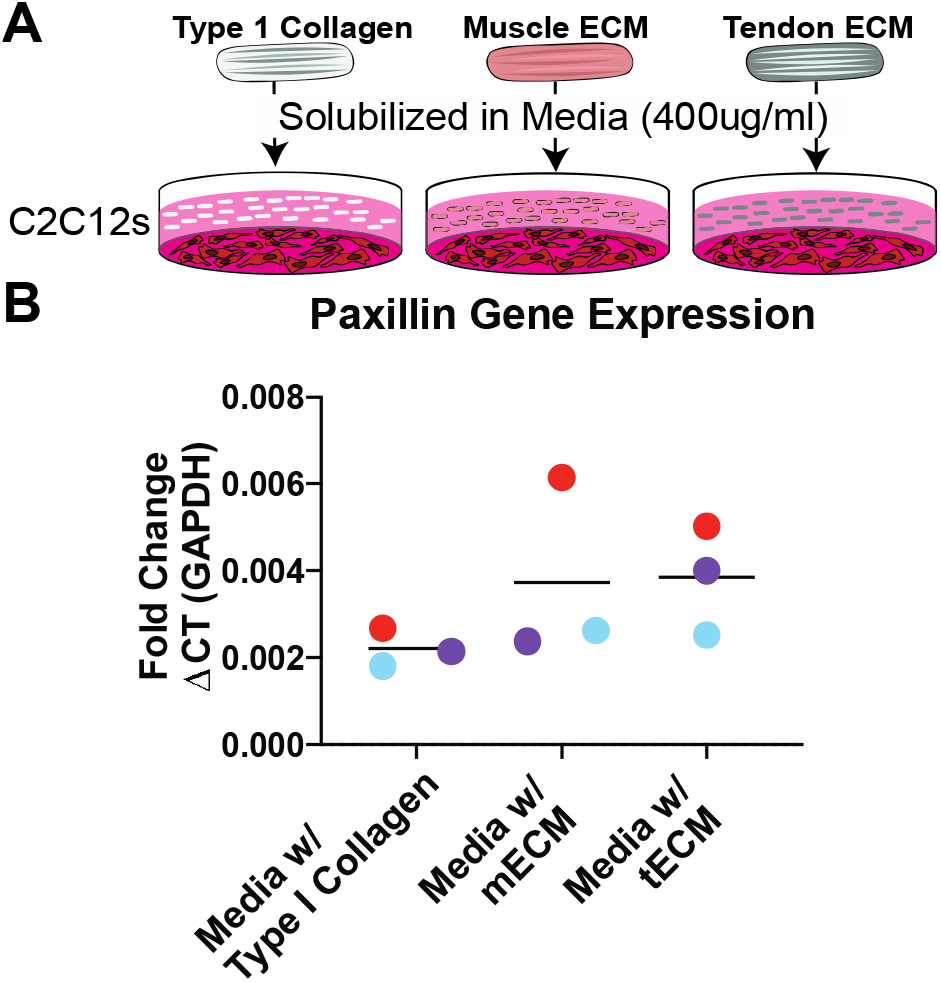
C2C12s cultured in ECM conditioned media did not have a noticeable effect on paxillin expression. A) C2C12s were seeded in cell culture plates, then cultured in proliferation media supplemented with type I collagen, mECM or tECM at a concentration of 400ug/ml of media. B) qPCR analysis did not show significant difference in paxillin expression between cells grown in ECM conditioned media and type I collagen conditioned media (p>0.05). ΔCT was normalized to housekeeping gene GAPDH, data is presented as fold change of ΔCT with the mean and individual data points of n=3 experimental replicates and colors representing paired replicates.

### 2.6 Paxillin and Type XXII Collagen Expression in Cells seeded in Tissue Specific ECM Tissue

Custom PDMS wells (15 mm x 10 mm x 4 mm) with two posts 8 mm apart and 2 mm in diameter were used to support 3D culture as tissues would contract around the posts. C2C12s and TFs were detached and suspended in either 5mg/ml type I collagen, 5mg/ml mECM or 5mg/ml tECM at 1×10^6^ cells/ml of gel. 5mg/ml was chosen as it is near the maximum concentration that is readily workable based on prior work from our lab after accounting for mixing with cells in media.^32^ The cell-laden ECM was seeded into the custom inserts to allow for culture. Gels were allowed to self-assemble for 30 min at 37C to fully form. After which proliferation media was used to culture tissues (Figure 6A). Tissues were cultured for 5 days, and media was changed at D3. At the conclusion of culture, the tissues were fixed and sucrose treated to improve sectioning. Tissues were cryo-sectioned at 10 μm. Samples were permeabilized, blocked, then treated with rabbit anti-paxillin antibody (Millipore-Sigma) at 1:100 dilution or rabbit anti-col22a1 antibody (ThermoFisher) at 1:200 dilution then both were subsequently treated with anti-rabbit donkey AlexaFluor 594 (Invitrogen) and sealed with Prolong Anti-Fade mountant with DAPI (ThermoFisher), the dilutions were suggested from the manufacturer. Anti-paxillin and DAPI were verified by staining TFs grown on tissue culture plastic and comparing the expected locations of staining with the cellular cytoskeleton (Supplemental Figure 2). To verify that the samples were being saturated with primary antibody (independent of the varying sample cross-sectional area) 2X and 4X concentrations of primary antibody was applied to C2C12s in type I collagen samples, which had the highest cross-sectional area (Supplemental Figure 3). For image analysis, 6 measurements per sample were used, more detail is described in Supplemental Figure 4. Images were acquired with an Olympus fluorescence microscope at 20X magnification. Parameters for image capture were consistent across all samples and different stains. Post-processing of brightness and contrast was consistent within staining groups. The ratio of TXRed and DAPI positive areas was defined as the Expression Index (E.I.) (Supplemental Figure 4). Percent contraction was determined using ImageJ to calculate the area of the tissue at the time of seeding and at 5 days, when the tissue had fully contracted (Supplemental Figure 5).

**Figure 6:**
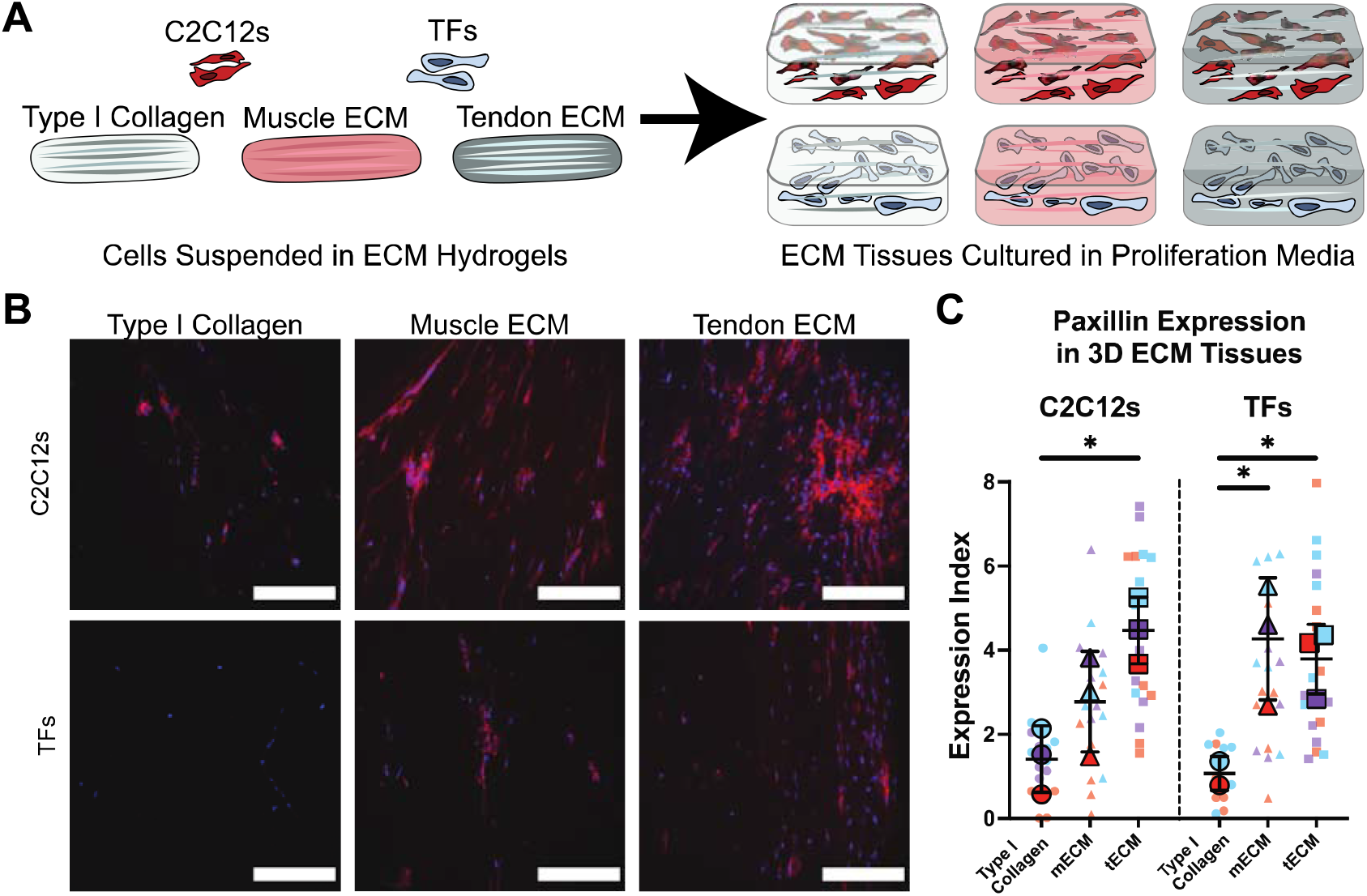
C2C12s and TFs seeded in 3D hydrogels had higher expression of paxillin in ECM tissues after being cultured for 5 days in proliferation media. A) C2C12s or TFs were suspended in type I collagen, mECM or tECM pre-gelation, then seeded in custom inserts for culture. Stable hydrogels were cultured in proliferation media for 5 days, before they were cryo-sectioned and stained. B) Representative fluorescent images of sections stained with anti-paxillin rabbit antibody and DAPI, showed that cells in ECM groups became more concentrated most likely due to contraction of the ECM tissue, where cells in type I collagen did not contract. Additionally, cells in ECM tissues had more positive staining for paxillin, shown in red here. Scale bars are 200 um. C) Paxillin protein expression was quantified using an expression index, which was defined as the positive area for TXRed channel normalized to the positive area for DAPI. Colors indicate measurements within the same tissue sample. Different colors represent different tissue samples. Mean values for each tissue sample are presented in the foreground with mean and SD for each condition. In C2C12s, tECM had the highest expression index, which was significantly different from type I collagen tissues. In TFs, both ECM tissues had significant increases compared to type I collagen tissues. Asterisks indicate a significant difference between ECM tissues and type I collagen (p<0.05), determined with one-way ANOVA (n=3 samples; 6 E.I. measurements per sample).

### 2.7 Paxillin Expression due to Contraction of ECM Tissue

Similar to ECM hydrogels for 3D culture, custom inserts were used to seed ECM hydrogels. C2C12s were seeded in 2mg/ml and 5mg/ml type I collagen (low-Col., high-Col. respectively) and 5mg/ml tECM at a cell density of 2×10^6^ cells/ml of gel. Tissue construct contraction was predicted in low-Col. and tECM (Figure 7A). Cells were cultured in proliferation media and analyzed in the same manner as ECM hydrogels for 3D culture.

**Figure 7:**
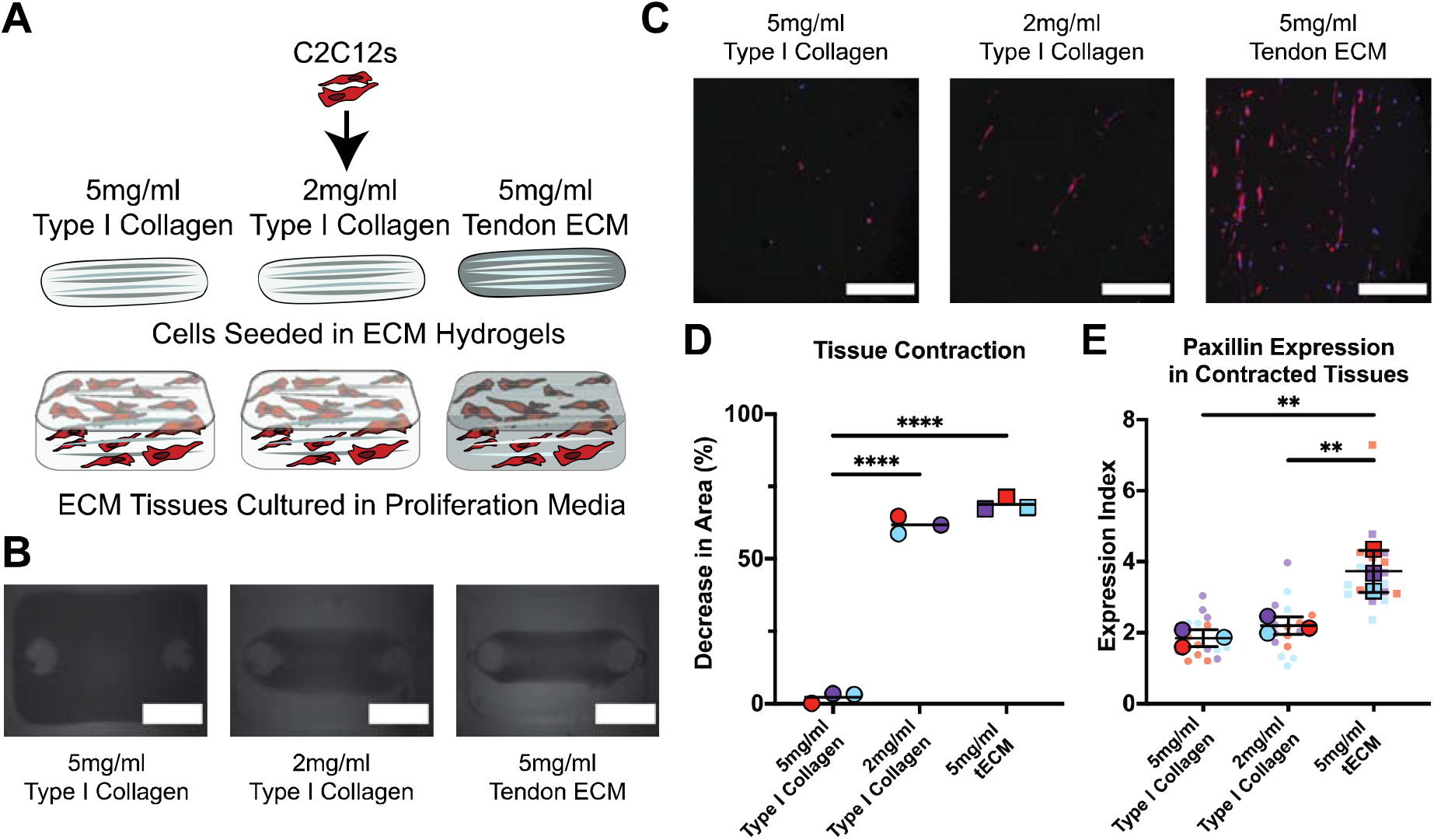
Contraction of hydrogel tissues did not induce paxillin expression. A) C2C12s were encapsulated in 2 concentrations, 5mg/ml and 2mg/ml, of type I collagen, and in 5mg/ml tECM. The formed tissues were cultured for 5 days in proliferation media before cryosectioning and staining. B) Brightfield images of tissues after 5 days of culture showed that decreasing the concentration of type I collagen allowed the cells to contract the tissue similar to tECM. Scale bars are 5 mm. C) Fluorescent images showing paxillin (red) and DAPI (blue) seemed to follow trends from other 3D tissues in that type I collagen did not have high paxillin expression, while tECM had higher expression. Scale bars are 200 um. D) Contraction was quantified by normalizing area of the contracted gels to the area of the seeded tissue after gelation, the mean contraction of 3 individual samples is shown here. The lower concentration of type I collagen and tECM both had higher contraction than the high concentration of type I collagen. Colors represent different ECM tissues. Four asterisks indicate a significant difference between contracted tissue constructs and 5mg/ml type I collagen constructs (p<0.0001) determined with one way ANOVA (n=3 samples). E) Paxillin expression was quantified with the expression index method previously mentioned. Both type I collagen groups had significantly lower expression of paxillin compared to tECM. While tissues made from at 2mg/ml type I collagen did contract, they did not have increased paxillin expression suggesting contraction does not increase this expression. Different colors represent different tissue samples, and E.I. measurements. Mean values for each sample is presented in the foreground, with mean and SD for each condition. Two asterisks indicate a significant difference between ECM tissues and type I collagen (p<0.01), determined with one-way ANOVA (n=3 samples; 6 E.I. measurements per sample).

### 2.8 Statistical Methods

To quantify decellularization, percent removal of DNA was calculated for decellularized tendon and muscle by normalizing amount of DNA in a sample prior and after decellularization. In ECM conditioned media experiments gene expression for Paxillin was measured with q-PCR in 3 different experimental samples per media condition. One-way ANOVA between media groups were paired by experimental replicates with Tukey post-hoc analysis. For 3D ECM tissues, n=3 biological samples were sectioned at 3 depths (TFs in type I collagen had n=2 biological samples, due to sample fracture during sectioning) and imaged in 2 areas (Supplemental Figure 4). Expression Index was measured 6 times for each biological sample (3 depths and 2 areas per depth) and averaged for n=3 biological replicates. One way ANOVA between tissue ECM was run using n=3 of biological replicate averages with Bonferroni’s multiple comparisons tests. Analysis was done separately for C2C12s and TFs. Similar to 3D ECM Tissues, E.I. measurements of 3 sections and 2 areas per section for each biological replicate were averaged for each biological replicate. To determine statistical significance in contraction experiments, one way ANOVA compared percent contraction in each ECM group which had a biological replicate of n=3. Similar to 3D ECM tissues, paxillin expression in contracted tissues was compared with one way ANOVA comparing the averages of n=3 biological replicates with Tukey’s multiple comparisons tests.

## 3. Results

### 3.1 Muscle and tendon can be decellularized to yield tissue specific hydrogels

Porcine muscle and tendon tissue (Figure 2A) were decellularized using the same decellularization protocol, and both were successfully digested to form hydrogels. Removal of DNA was 98% for muscle tissue and 96% for tendon tissue, determined with PicoGreen DNA quantification assays (Figure 2B). Additionally, H & E staining of tissue showed removal of cell nuclei (Figure 2C) confirming DNA removal and retention of some ultrastructural features, like collagen crimp in decellularized tendon. The gross tissue structure of decellularized ECM is disrupted, likely due to freezing and thawing of tissue before decellularization. While this may be a concern in other applications, the ECM is digested completely for use in a hydrogel so native macrostructure does not need to be maintained. In addition, ECM tissues and type I collagen were stained for DAPI, anti-paxillin and anti-Col22a1 to verify removal of nuclear content and that there was no presence of the paxillin or type XXII collagen within the matrix, all of the tissues were negative for those proteins (Supplemental Figure 1).

### 3.2 Proteome of ECM Materials

The proteomic discovery identified around 2,200 proteins for mECM and 1,900 for tECM, with 570 proteins uniquely detected in mECM in contrast to 238 for tECM. Focusing our GO analysis on ECM-associated proteins and subunits, the pie charts in Figure 3 show that the composition (by number of proteins) of the two materials are close to each other. It’s important to note that this represents the number of proteins identified as ECM-associated and does not represent tissue composition by mass. However, there were differences in the abundance of the proteins under this GO-ECM classification, their abundances based on the Log2 (LFQ tECM/mECM) can be found in Supplemental Figure 6.

### 3.3 Rheological Properties were similar for mECM and tECM Hydrogels

Rheology was used to analyze differences in the mechanical properties of the hydrogels produced from the different materials. Type I collagen at a concentration of 5mg/mL was found to have a higher complex viscosity with a mean of 7.798 Pa*s across the frequency sweep from 100-26 rad/s (Figure 4). This was significantly higher than the 2mg/mL type I collagen (0.6652 Pa*s), tECM (0.7993 Pa*s), and mECM (2.019 Pa*s) across the same frequency sweep (p<0.0001 for each comparison) (Figure 4). The mECM was also significantly higher than the 2mg/mL type I collagen and the tECM (p<0.001 and p<0.01 respectively). The tECM and 2mg/mL type I collagen were not statistically different.

### 3.4 ECM conditioned media does not promote paxillin

To assess bioactivity of the digested ECM materials independent of any structural cues, C2C12s were cultured in media supplemented with digested tissue specific ECM (Figure 5A). Relative to controls, mean paxillin expression increased by 65% and 74% in mECM and tECM conditioned media, respectively (Figure 5B). Relative paxillin gene expression had high variability between biological replicates in each of the ECM supplementation groups, and there was no statistically significant effect of media supplementation (p=0.19).

### 3.5 3D ECM hydrogels promoted paxillin expression

To create a more physiologic microenvironment, C2C12s and TFs were embedded within 3D hydrogels made from tissue specific ECMs (Figure 6A). Cells cultured in 3D tissues had different levels of gel contraction after 5 days of culture (Supplemental Figure 5) resulting from contractile forces of the cells in the matrix expelling water from the hydrogel. C2C12s and TFs in type I collagen contracted least of all the tissue groups with little to no noticeable decrease in surface area. mECM tissues had a more variable decrease in area, while tECM consistently contracted.

Via fluorescence imaging, C2C12s had the highest paxillin expression when cultured in the tendon ECM (Figure 6B). TFs also had higher expression in tissue derived ECM than when cultured in type I collagen. Furthermore, contraction seemed to increase cell density, which is expected contracted gels have reduced volume for the same number of seeded cells. In type I collagen, both cell types had some paxillin expression, whereas in other groups, especially C2C12s in tECM, paxillin expression seemed to be upregulated all around the cell and throughout the matrix. It was verified that paxillin was not present in any of the ECM materials used for the 3D tissues. Finally, in tissues with higher levels of paxillin expression, the highest level of positive staining seemed to occur in areas where more cells were in contact with each other.

Fluorescence images were quantified using a ratio of positive signal area of each channel that we define as E.I. (Supplemental Figure 4). Within each sample, measurements at different depths and areas had similar ranges of values, showing that that staining wasn’t concentrated in different parts of the ECM tissues. One way ANOVA was used to compare these differences across the 3 biological replicates. Similar to quantitative images, there is an increase in expression in ECM tissues compared to type I collagen (Figure 6C). The highest E.I. occurred in C2C12s in tECM, with a mean 68% higher than C2C12s cultured in type I collagen (p<0.05). C2C12s in mECM had a mean increase of 48% over those cultured in type I collagen (p>0.05). TFs exhibited similar expression patterns, with the mean E.I in mECM and tECM tissues being 75% and 71% higher, respectively, than in type I collagen In that comparison, both groups had a significant increase over type I collagen at p<0.05. In contrast to supplementing media with ECM, culturing C2C12s and TFs in tissue specific ECM caused an up regulation of paxillin expression in both cell types.

### 3.6 Contraction of tissues does not lead to paxillin expression, in type I collagen constructs

In previous experiments, tissue construct contraction was dependent on the ECM material (Supplemental Figure 5). Independent of what cells were used in the tissue constructs, tECM constructs had the most contraction, while type I collagen constructs contracted the least. To verify that varying levels of gel contraction did not have an effect on paxillin expression, C2C12s were cultured in 2 different concentrations of low-Col. (2 mg/ml) and high-Col. (5 mg/ml), yielding a non-contracting tissue construct and a contracting tissue construct respectively as well as tECM, which also yielded a contracting tissue construct.

C2C12s were cultured for a period of 5 days in proliferation media in each of the three tissue environments. Representative images are shown at day 5 of culture in Figure 7B. There is a clear decrease in surface area in the low-Col. tissue and in the tECM tissue group while the high-Col. tissue does not seem to have any change from the initial area of the hydrogel at the time of seeding. This decrease was quantified using ImageJ, by measuring the outline of the tissue construct at the end of the 5 day culture period. Overall, the high-Col. tissues had a mean decrease in area of only 3%, while the contracted tissues low-Col. and tECM had decreases in area of 62% and 68%, respectively (Figure 7D). The contracted tissues were significantly more contracted than high-Col. (p<0.0001).

Representative fluorescent images are shown in Figure 7C. Similar to previous findings, high-Col. did not have a lot of positive staining for paxillin, while the tECM had clear paxillin expression, corroborating the results from 3D ECM tissues, even with the increased cell density. Interestingly, low-Col. did not have an increase in positive staining compared to high-Col., even though the tissue contracted similarly to tECM. Paxillin expression was quantified with image quantification methods previously described. Overall tECM had double the expression of the type I collagen groups. One-way ANOVA was used for analysis of the 3 biological replicates (each with 6 measurements for EI), in which tECM had a statistically significant increase in E.I. of paxillin (p<0.01).

### 3.7 Type XXII collagen is present in 3D ECM tissues

3D ECM tissues were prepared and cultured for 5 days using the same protocol for 3D tissues used to study paxillin expression (Figure 8A). The tissues had similar contraction to the previously described tissues. E.I. was quantified the same way as previously described in paxillin expression studies. Fluorescence images of 3D ECM tissues, shows positive staining for type XXII collagen (Figure 8A). Immunohistochemical analysis of type XXII collagen showed similar results to paxillin expression in 3D ECM tissues, however the relative differences between groups were not as great, quantitatively. The most type XXII collagen staining was observed in tECM for both cell types. While there was not a statistically significant increase between C2C12s in tECM, mean E.I.=2.1, compared to type I collagen, mean E.I.=1.5, (p=0.47), TFs had significantly higher expression of type XXII collagen (p<0.05), the expression index of TFs in tECM was about two times that of TFs in type I collagen (Figure 8B). In mECM tissues, an increases in type XXII collagen expression relative to type I collagen is not evident after 5 days of culture.

**Figure 8:**
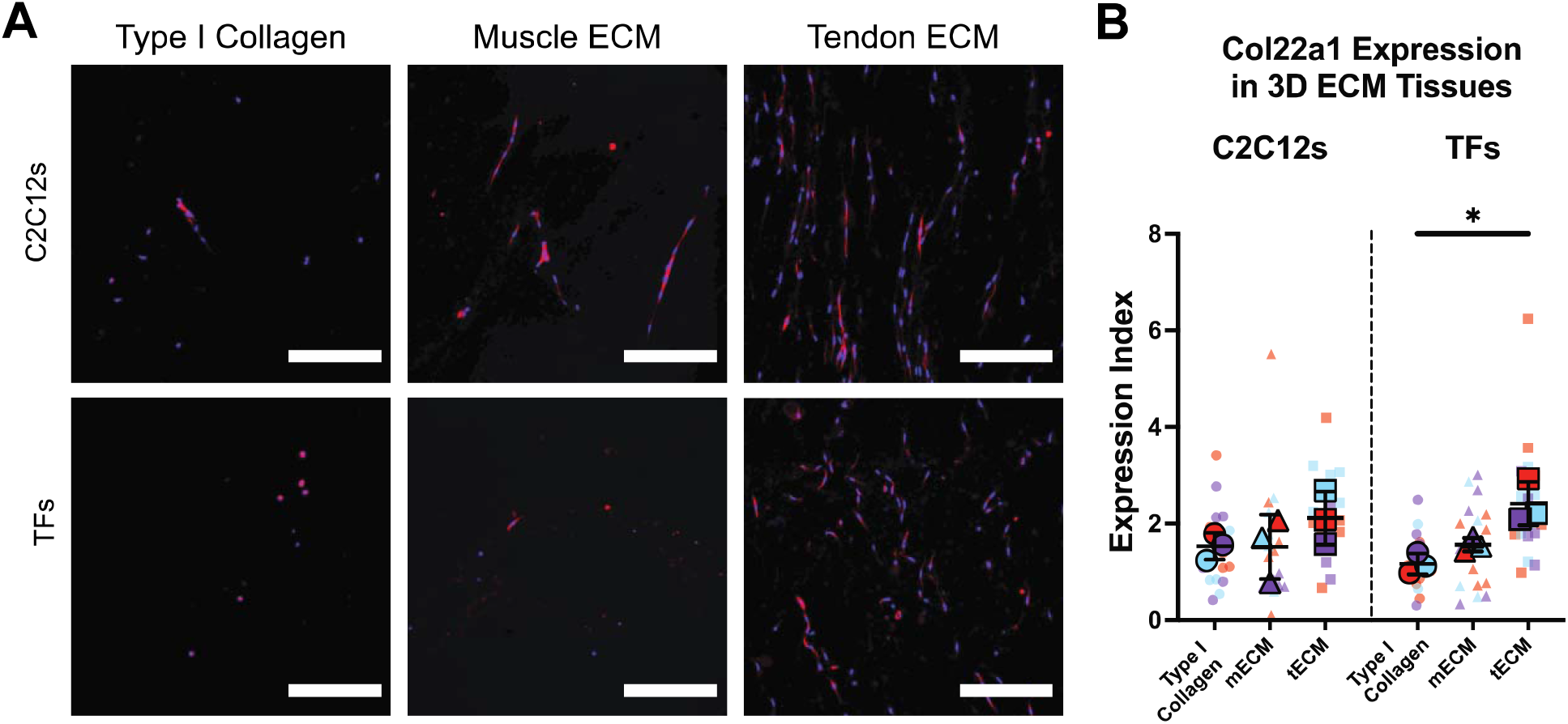
A) Representative fluorescent images of sections stained with anti-col22a1 rabbit antibody and DAPI scale bars are 200 um. Cells cultured in tECM tissues have more positive staining for type XXII collagen. Cells in mECM constructs had similar expression to cells cultured in type I collagen constructs. B) Type XXII collagen protein expression was quantified using E.I. colors indicate measurements within the same tissue sample. Different colors represent different tissue samples, n=3. Mean values for each tissue sample are presented in the foreground with mean and SD for each condition. Compared to TFs in type I collagen, TFs in tECM had the largest relative difference, which was statistically significant (p<0.05). Expression levels were similar to paxillin expression patterns, however the relative differences between groups were not as pronounced.

## 4. Discussion

The myotendinous junction consists of unique ECM, structural features, cell-matrix and cell-cell interactions.^10^ Understanding how the unique aspects of the junction are formed and how they are maintained are important to inform regenerative medicine strategies to alleviate loss of tissue function in the event of disorders or traumatic injury. In this study, we hypothesized that myoblasts and tenocytes would be influenced by a tissue-specific microenvironment, resulting in a junction-like cellular response. In the present study, we focused on the expression of a junction-like protein paxillin, as well as a more specific extracellular matrix protein type XXII collagen, and the use of muscle and tendon derived extracellular matrix hydrogels to mimic the composition of the respective tissue microenvironments. Cells cultured in monolayer on tissue culture plastic with media supplemented with tissue-specific ECM did not show increased paxillin protein expression compared to type I collagen supplemented media. However, C2C12 and TF cells cultured within ECM hydrogels showed increased paxillin protein expression compared to type I collagen. Type XXII collagen was expressed more in samples cultured in tECM hydrogels, but the relative increase to type I collagen was not as high. Even though these ECM constructs contracted over time, the contraction had no effect on paxillin expression, supporting the hypothesis that ECM-specific components were the driving force behind the increase in paxillin expression. The *in vitro* system presented here offers a unique method to study cell-matrix interactions by deconstructing the junction to study how cells interact with opposite matrix and how that may drive junction specific cell attachment to the matrix.

Tissue specific ECMs have emerged as a unique biomaterial class that promotes regenerative phenotypes in immune cells and attracts stem cells as it is broken down.^17^ Tissue specific ECM derived from muscle and tendon has been used in previous *in vitro* and *in vivo* studies^20–25^ but junction specific interactions remain under-explored. Here we use the ECM from muscle and tendon together to determine the effects of tissue specific cell-matrix interactions to model the junction where cells of one tissue may be in contact with matrix of the other tissue. These materials were derived from porcine Achilles tendon and gastrocnemius by excising muscle tendon units and then separating the two tissues. While many labs use tissuespecific decellularization protocols, this study used the same process for both musculoskeletal tissues allowing for direct comparison between the inherent signals present within each tissue-specific ECM. The protocol modified from Wolf et al^22^ was effective for both tissues, with muscle and tendon ECM having about 3%-4% of the original DNA content remaining. This removal was confirmed with hematoxylin and eosin staining before and after the decellularization protocol.

Proteomic analysis of the two ECM derived materials revealed that after the decellularization, they contain many different ECM related proteins. While the general ECM makeup of each material was similar categorically, the individual proteins had varying levels of abundance when comparing mECM with tECM (Supplemental Figure 6). Previous protoeomic analysis of mouse ECM across the MTJ by Jacobson, et. al. 2020 characterized specific ECM proteins present in muscle, tendon or at the MTJ junction.^11^ Some muscle specific ECM proteins identified by that analysis (LAMA2, LAMB1, NID1, TGM2) were also present in higher relative abundance in the mECM material in the current study. Tendon specific ECM proteins identified in the prior work (COL12A1, CHAD, CILP2, COMP, TNC, VCAN, THBS1, THBS2, MFGE8, SERPINF1, ABI3BP, COL2A1 and LOX) were also found in relative higher abundance in the tECM in the current work. However, there were a few discrepancies between the proteome of the ECM materials and those previously reported for muscle (LAMC1, NID2, LGALS1, COL4A1 and LAMB2) and tendon (POSTN and FGG), which is interesting for future characterization of the ECM materials. These discrepancies most likely arise from the decellularization process, where there is likely some removal of ECM proteins as the cellular content is removed. However, it is interesting that the tissue specific materials do maintain a vast majority of the specific ECM proteins that Jacobson et. al. identified in their thorough analysis, and that the remaining ECM proteins could elicit production of MTJ-specific proteins from the embedded cells in this study. The discovery proteomics used to analyze the two materials is a preliminary look at the inherent differences between the ECM used. More targeted analysis, especially of proteins that could influence the cellular responses in the materials, is needed to fully characterize the differences. Further characterization could be used to link the proteome of the tissue specific ECM to cell responses as well as comparing the tissue specific properties maintained after decellularization.

Tissue specific ECM has been used in several forms including lyophilized sheets, powders to supplement media and in a hydrogel form.^33^ Tissue specific ECM in solubilized form as a media supplement can influence immune cells^31^ and stem cells.^34^ To determine if solubilized ECM affected C2C12s, solubilized mECM and tECM was added to proliferation media during monolayer culture on tissue culture plastic. The solubilized matrix did not change the expression of paxillin when compared to type I collagen supplementation. The lower overall concentration of tissue specific material in the supplemented media could be below a threshold needed to illicit a cell response. This was the upper limit for supplementing media with muscle and tendon ECM as higher concentrations resulted in gelation of the media. Additionally, myoblast cells may not express MTJ specific integrins when attached to tissue culture plastic in monolayer. This is consistent with observations by Merceron et.al., where myoblasts and fibroblasts in contact with synthetic materials did not form native focal adhesions without being in contact with natural ECM.^13^ Determining if the significant paxillin expression occurs because of differences in cell morphology in a 3D environment or how the ECM components are presented to the cells is worthy of exploration in future studies.

ECM hydrogels allow cells to be suspended within the matrix, which creates a more relevant microenvironment for the cells to attach and spread when compared to monolayer culture. The cell-laden hydrogels had varying levels of contraction with tECM tissues having the largest decrease in surface area. Paxillin was expressed more in myoblast cells seeded in tECM compared to type I collagen. There was also more expression in cells seeded in mECM, albeit not as much of an increase than in tECM. Similarly, paxillin was expressed more in tenocytes when cultured in tissue specific matrix when compared to type I collagen. While 3D culture of cells in hydrogel tissues may have up-regulated paxillin expression, the increased expression was not as evident when cells were cultured in 3D type I collagen hydrogels. Similarly, type XXII collagen was visualized using IHC with the most expression found in tECM hydrogels. The relative differences between cell and ECM conditions were similar but not as large to those observed with paxillin expression. IHC showed that type XXII collagen is being expressed by the cells in tECM constructs especially, however larger expression may occur outside of the 5 day timeframe used in the present study. This suggests that cells cultured in tissue specific ECM interacted with the matrix differently than in type I collagen. This finding supports the notion that our model can produce integrin-associated, and MTJ extracellular matrix protein expression in myoblasts and tendon fibroblasts that is known to be expressed at the myotendinous junction. Additionally, there was differential expression of the same cells in one matrix compared to another. Paxillin is part of an integrin mediated complex that anchors muscle cells to tendon ECM. Within muscle tissue, the primary complex is the dystrophin associated complex. We believe that C2C12s respond differently to tendon ECM (tECM) because the tECM promotes the integrin mediated complex. This result was exciting because it suggests that this in-vitro system can potentially be used to capture a phenomenon observed in native tissues. This exemplifies an interesting characteristic of these tissue constructs, the ECM hydrogels promoted MTJ integrin associated proteins and it can be determined which cell-matrix interaction had the greatest up-regulation. This is useful information because we can begin to dissect specific interactions that may contribute to the development of the junction. It also supports inclusion of a native-like MTJ in 3D tissue models to study the function, development, and maintenance of muscle-tendon units.

The varying levels of contraction of the 3D engineered tissue presented a unique challenge, as the change in boundary conditions as well as the mechanical properties of the different hydrogels, could influence cellular responses. Interestingly, the changes in complex viscosity of the hydrogels were inversely related to the contraction of each material in the tissue constructs. ECM hydrogels had lower complex viscosity and showed more contraction than type I collagen at 5 mg/ml. C2C12s in tECM had the largest difference in paxillin expression compared to type I collagen, as well as a large amount of tissue contraction. To verify that paxillin expression was not a result of the mechanical environment in contracted tissues, the C2C12-laden hydrogels were adjusted to allow contraction of type I collagen tissues that resembled the contraction of tECM tissues. A lower type I collagen concentration of 2 mg/ml was used to allow similar levels of complex viscosity and contraction as the tECM at 5 mg/ml. However, expression of paxillin in the 2mg/mL type I collagen constructs was similar to 5mg/mL type I collagen constructs, that did not contract. While direct comparison with Freytes et. al. is difficult because of different materials and concentrations, complex viscosity values are within a similar range with lower concentrations of type I collagen behaving like higher concentrations of ECM hydrogels. In addition, hydrogel concentration had a direct effect on complex viscosity. With the described methods used to form the ECM hydrogels, the 5 mg/ml concentration used is at the upper limit after gel neutralization and buffering, and addition of the cell suspension. As such, it is not possible to make a tECM hydrogel that mimics the rheological properties of 5 mg/ml type I collagen. We cannot be certain how changes in mechanical properties within the tECM constructs would alter paxillin expression. However, we can conclude that for type I collagen, degree of contraction combined with varying mechanical properties did not influence observed paxillin expression.

This study is not without limitations. The primary marker that was focused on is known to be present at the myotendinous junction but is also present in many other tissues and cell types. Additionally, this study is focused on cell types in isolation from each other, without any cell-cell interactions. Protein analysis of the ECM of muscle-tendon units revealed that there are several proteins, specifically type V α3 collagen, and type XXII collagen, that occur primarily at the junction.^10,11^ Research into this specific protein synthesis could inform strategies for development of native-like muscle-tendon interfaces, increasing the function and efficacy of engineered muscle-tendon units. Future studies should include analysis of additional MTJ specific proteins like type XXII collagen, over longer time periods, along with cell signaling pathways leading to increased focal adhesion expression. De-construction of the muscle tendon unit may provide some insight to specific proteins and features of the MTJ; however, we are only examining individual cell types interacting with a single matrix. An accurate model of the muscle tendon unit should include incorporation of both cell types, interacting in their own tissue environments as well as with the opposite tissue environment. Incorporating the information obtained here in a multi-tissue model will allow for the study of the muscle tendon unit with a junction or interface component. This should be considered an important inclusion for muscle tendon unit models as proteins such as paxillin are up-regulated natively with mechanical loading, and lack thereof negatively affects the integrity of the junction.^12^ This research plays a vital role in determining composition of an interface region that can be used within a more complex multi-tissue model.

## 5. Conclusion

Overall, the objective of this study was to determine how cell-matrix interactions play a role in regulating paxillin expression by myoblast cells. This unique approach to modelling the MTJ allowed for cells of one tissue to be in contact with matrix from the other tissue which could provide clues to how matrix of opposite tissues influence development and maintenance of the junction. Tissue specific materials did modulate the expression of paxillin when used to culture cells in 3D. This finding is useful information for the development of a muscle tendon unit *in vitro* model that incorporates the two tissue types and a junction between them. This can be used to study interface specific characteristics, in tissues beyond muscle and tendon, as well as aiding in the formation of multi-tissue models with an interface region.

## Acknowledgements

The authors would like to acknowledge Amber Detwiler for her work on the contraction analysis of the tissues. Proteomics work was performed by the Molecular Education, Technology and Research Innovation Center (METRIC) at NC State University, which is supported by the State of North Carolina.

## Author Contributions

LG wrote the manuscript, planned and executed the studies presented. MF and DF wrote the manuscript, planned experiments and oversaw execution of the studies presented. ZD performed rheological analysis of ECM hydrogels and assisted in writing the manuscript. CM performed proteomic analysis and assisted in writing the manuscript.

