## Supplemental for "Extracellular Matrix Hydrogels Promote Expression of Muscle-Tendon Junction Proteins"

**Supplemental Data**

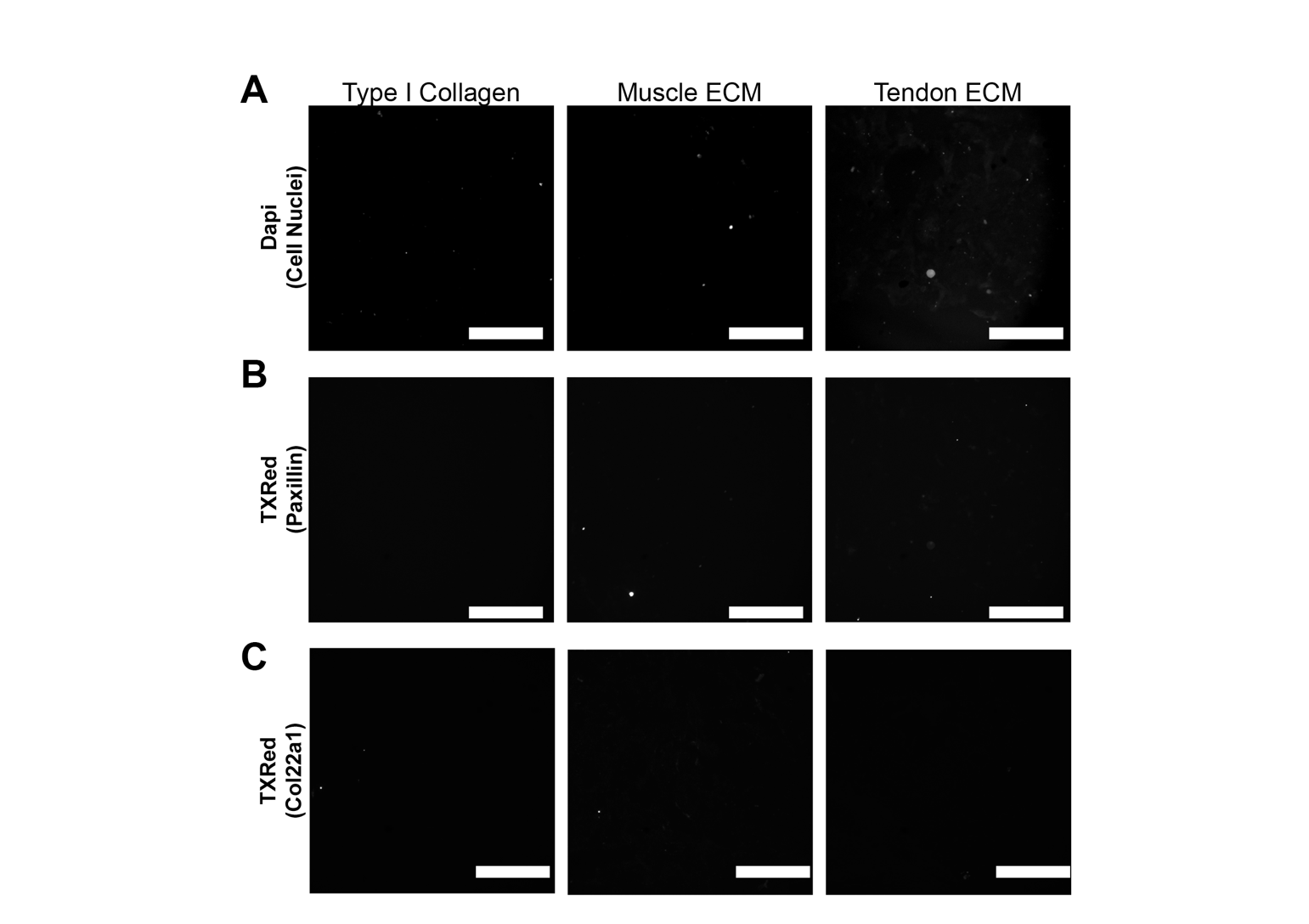

**Supplemental Figure 1**: ECM hydrogels and type I collagen were stained for A) Dapi, B) anti-paxillin and C) anti-Col22a1 to verify removal of nuclear content and that ECM gels were not positive for the proteins of interest. Images are presented in gray scale, the scale bar is 200 um.

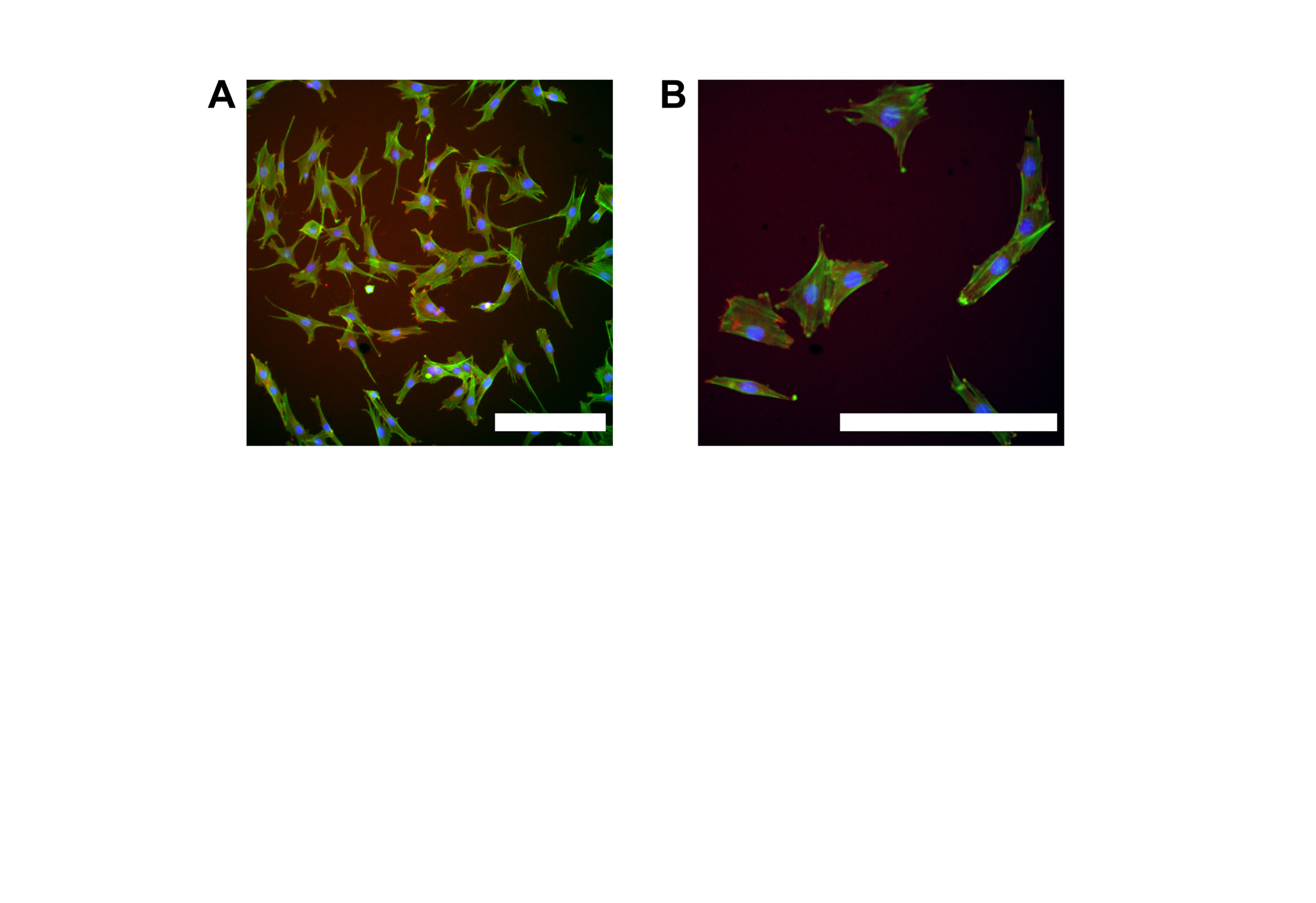

**Supplemental Figure 2:** A) 20X and B) 40X fluorescent images of tendon fibroblasts grown on tissue culture plastic and stained with DAPI (blue), anti-paxillin (red) and actin-488 (green). Scale bars are 200 um. This figure shows localization of DAPI within the actin cytoskeleton and the expression of paxillin at the edges of the cellular extensions, where focal adhesions would be expected.

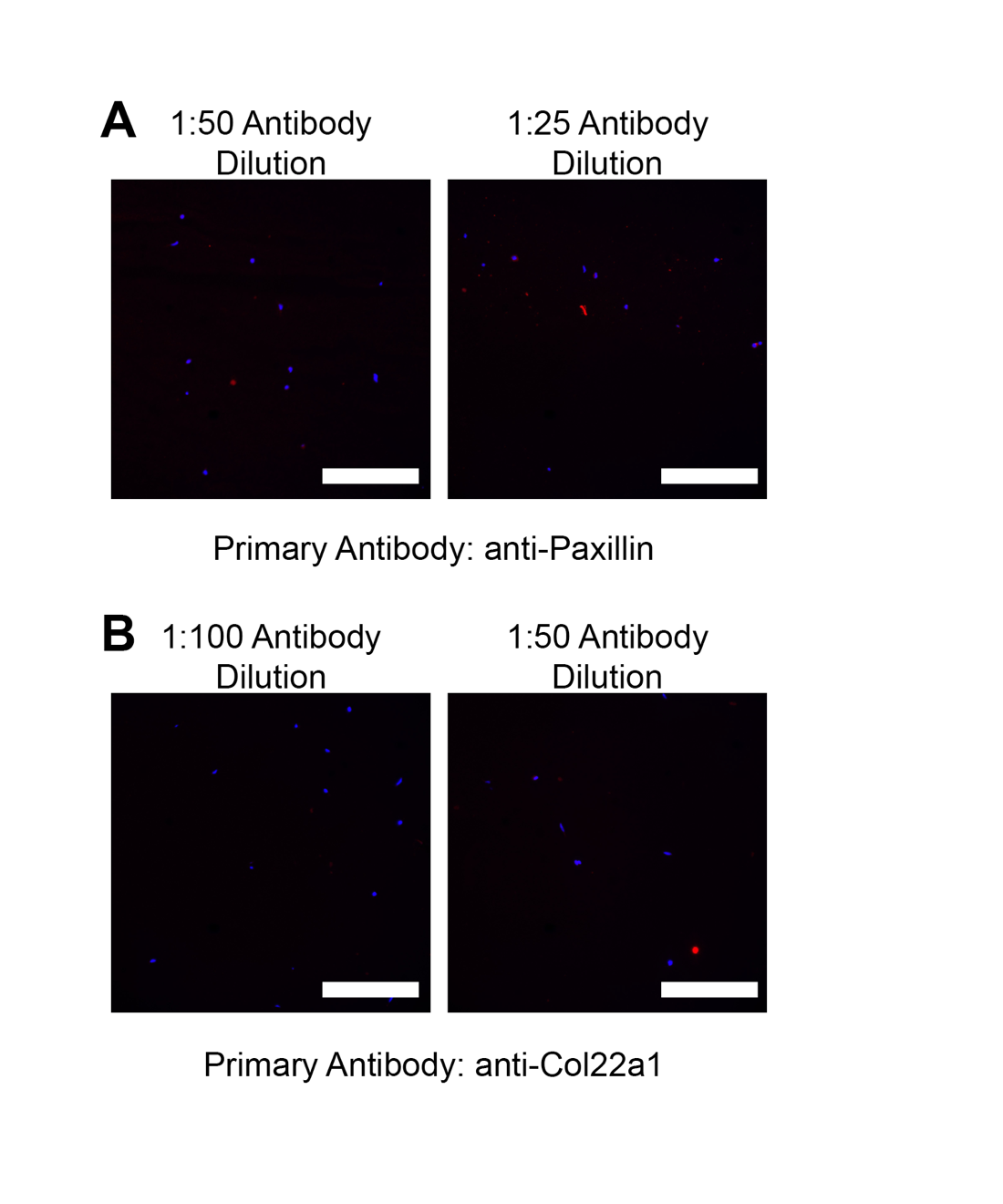

**Supplemental Figure 3:** To ensure that antibody availability was not a factor in varying stains for samples with larger cross-sectional area, C2C12s in type I collagen samples were stained as described in other experiments, except using 2X and 4X the recommended concentrations of 1:100 for anti-paxillin, and 1:200 for anti-col22a1. Scale bars are 200um. There is no difference between higher concentrations.

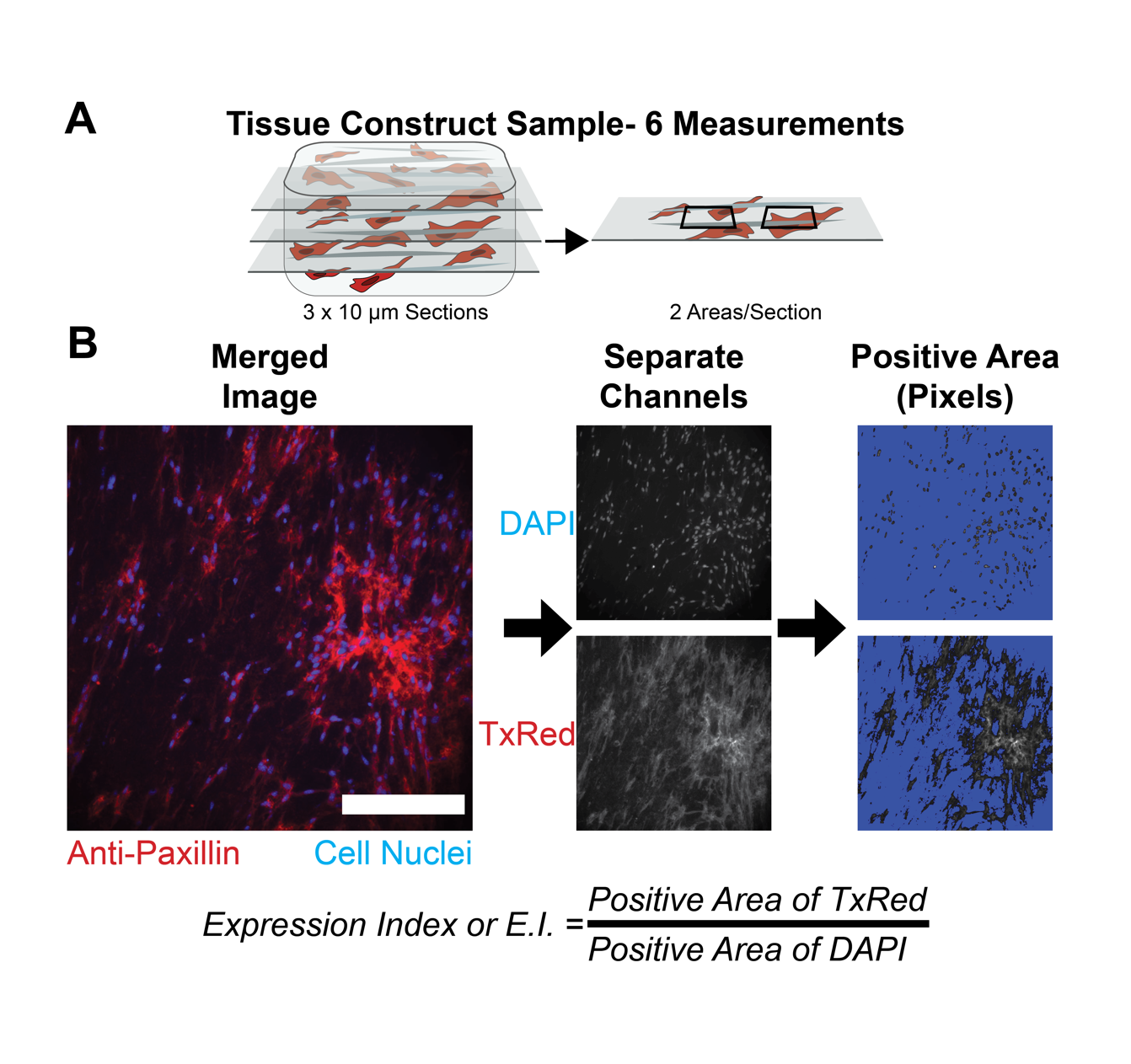

**Supplemental Figure 4:** A) Tissue construct samples were sectioned at 3 depths for immunohistochemistry. Each depth was imaged in 2 separate areas, so for each tissue construct 6 images were obtained to quantify paxillin expression. B) To quantify protein expression of paxillin, an expression index (E.I.) was calculated for samples. E.I. was defined as the positive pixels of the TxRed channel divided by the positive pixels of the DAPI channel. The higher the protein expression of paxillin (visualized with Anti-Paxillin antibody), the more positive area of the image. To factor in different cell densities within hydrogel tissues, the positive area of protein expression was normalized by the positive area of cell nuclei. The ratio of the positive areas should correlate if cells are expressing paxillin equally, however this ratio increased in tissue specific hydrogels.

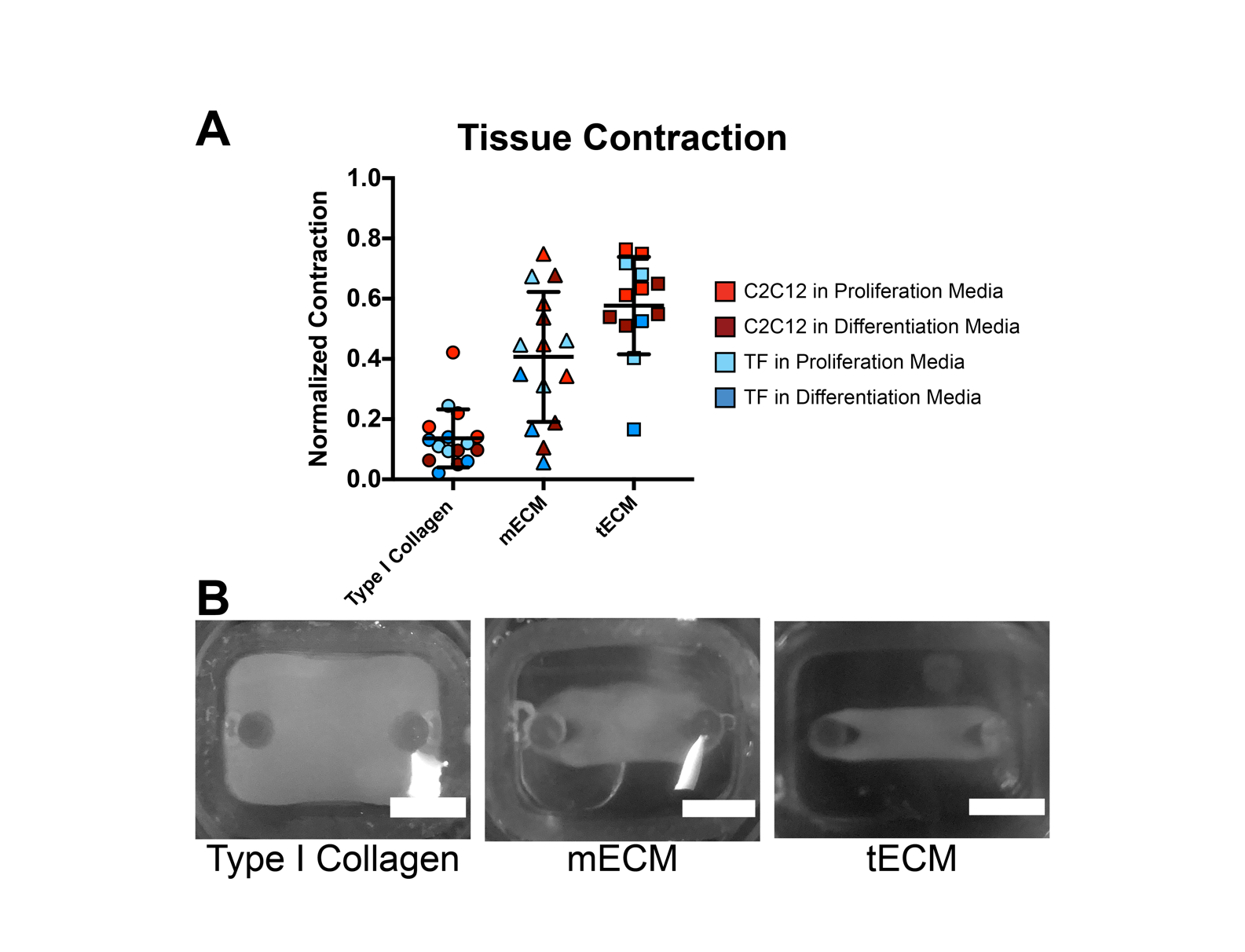

**Supplemental Figure 5:** After 48 hours cell-laden hydrogels contracted (C2C12 seeded hydrogels shown here). Interestingly, depending on the hydrogel, there was varying contraction amounts. All hydrogels were at an initial concentration of 5mg/ml. The trend of contraction, with type I collagen contracting the least and tECM contracting the most was consistent across cell types and media groups, represented by the same color in the plot. Cells in mECM and tECM contracted significantly more than cells in type I collagen (p<0.0001) determined with one way ANOVA comparing all cells in each matrix (n=16 samples). Varying contraction could be due to cells interacting differently with the microenvironment. The contraction increased cell density within the construct and likely affected the mechanical properties of the gel, although there was no analysis to confirm this.

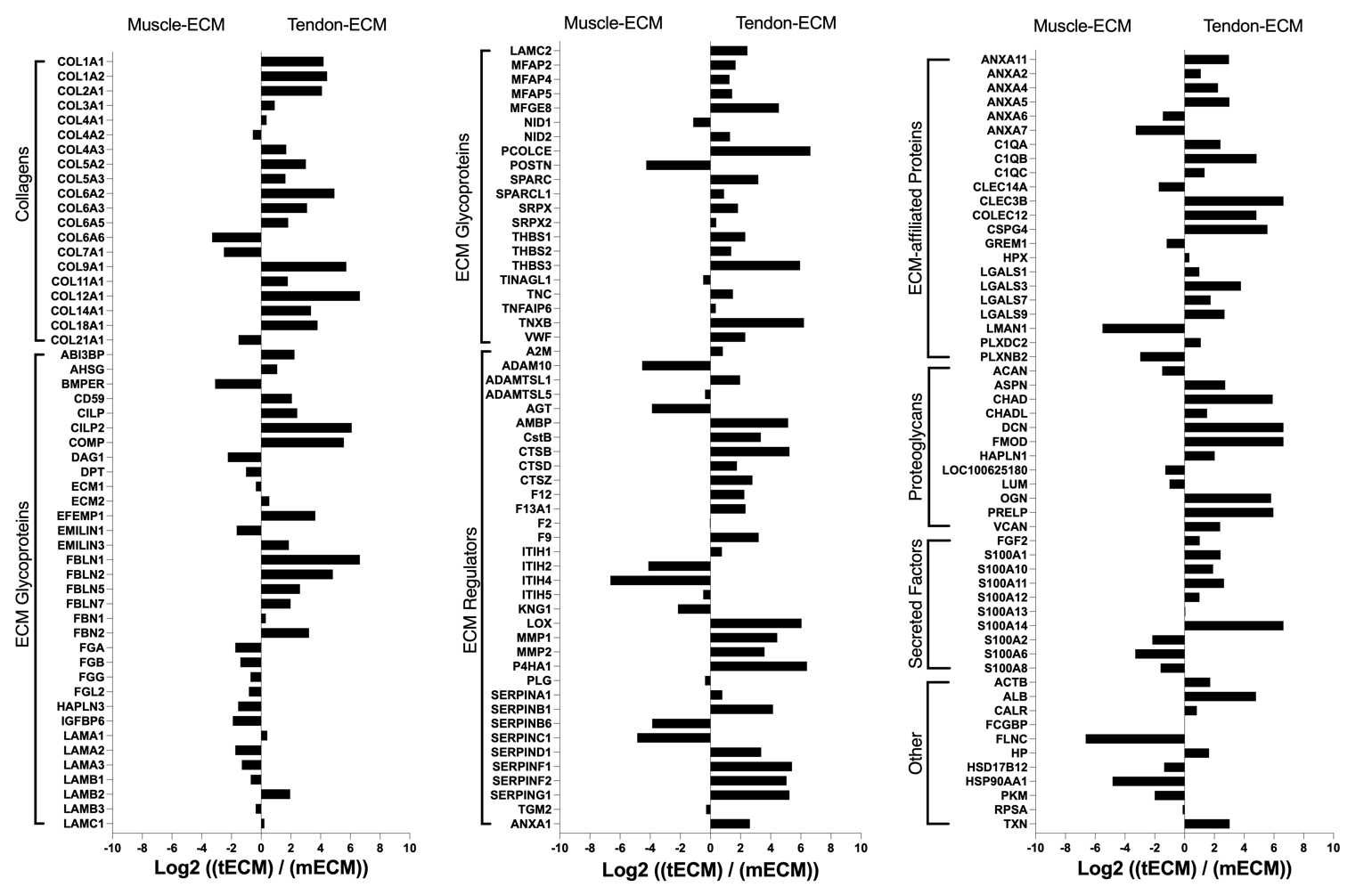

**Supplemental Figure 6**: Fold change of abundance of ECM-related proteins in tendon and muscle. Negative log change indicates an increased abundance of the protein in muscle ECM, while a positive value indicates and increased abundance of the protein in tendon. While overall, by protein category the materials are similar, there are many unique proteins within each category.

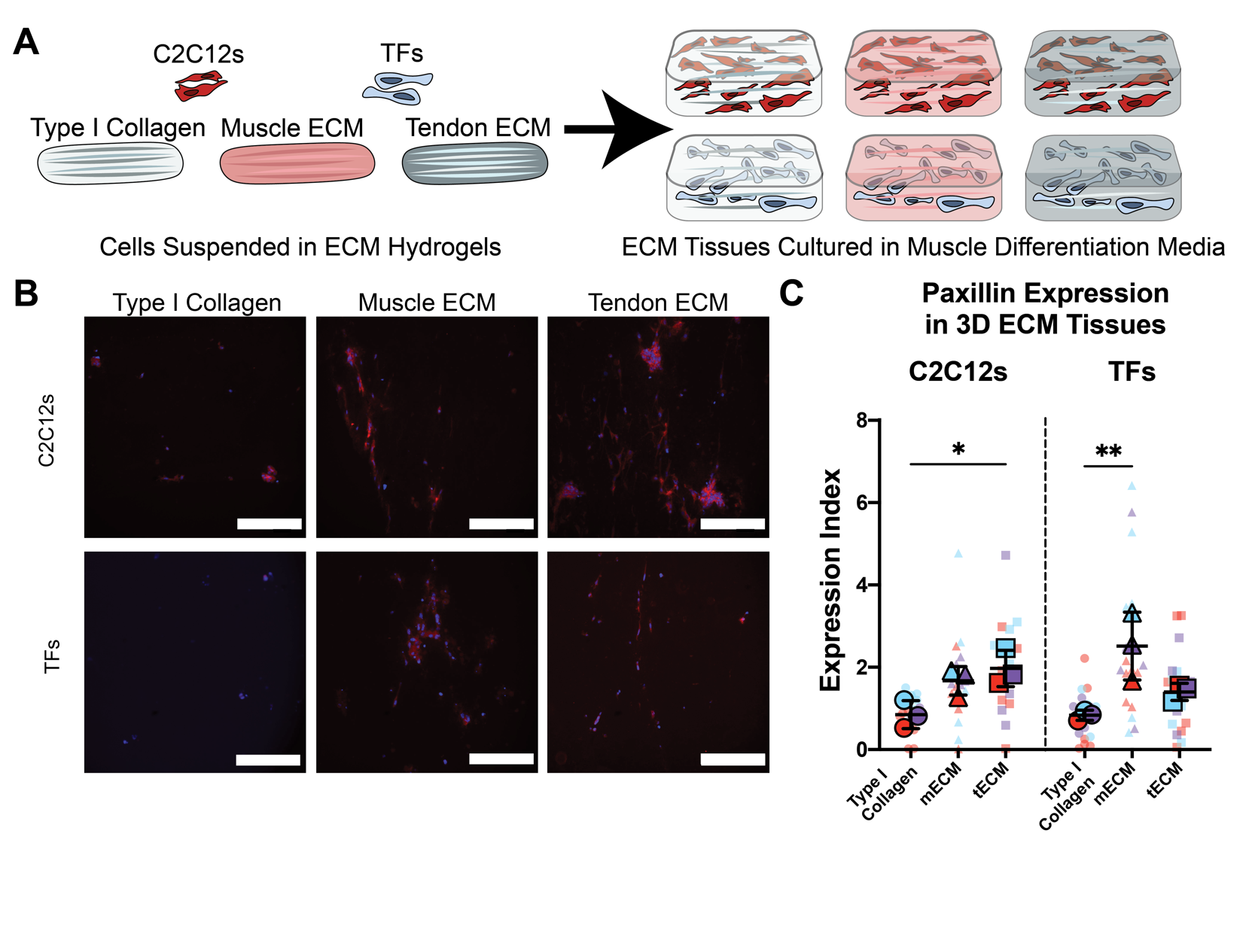

**Supplemental Figure 7:** 3D hydrogel tissues were also cultured in muscle differentiation media (mDiff). A) C2C12s or TFs were suspended in type I collagen, mECM or tECM pre-gelation, then seeded in custom inserts for culture. Stable hydrogels were cultured in mDiff for 5 days, before they were cryo-sectioned and stained. B) Representative fluorescent images of sections stained with anti-paxillin rabbit antibody and DAPI. Cells in ECM tissues had more positive staining for paxillin, shown in red here. Scale bars are 200 um. C) Paxillin protein expression was quantified using an expression index, which was defined as the positive area for TXRed channel normalized to the positive area for DAPI. Colors indicate measurements within the same tissue sample. Different colors represent different tissue samples. Mean values for each tissue sample are presented in the foreground with mean and SD for each condition. In C2C12s, tECM had the highest expression index, which was significantly different from type I collagen tissues. In TFs, mECM tissues had significant increases compared to type I collagen tissues. Asterisks indicate a significant difference between ECM tissues and type I collagen (* is p<0.05 and ** is p<0.01), determined with one-way ANOVA (n=3 samples; 6 E.I. measurements per sample). Overall trends in tissue constructs cultured in mDiff were similar to those cultured in proliferation media, but expression was not as high across all conditions.
